# Ormdl3 regulation of specific ceramides is dispensable for β-cell function and glucose homeostasis under obesogenic conditions

**DOI:** 10.1101/2023.02.11.528130

**Authors:** Liam D Hurley, Hugo Lee, Gina Wade, Judith Simcox, Feyza Engin

## Abstract

Chronic elevation of sphingolipids contributes to β-cell failure. ORMDL3 has been identified as a key regulator of sphingolipid homeostasis, however, its function in pancreatic β-cell pathophysiology remains unclear. Here, we generated a mouse model lacking *Ormdl3* within pancreatic β-cells (*Ormdl3*^β-/-^). We show that loss of β-cell *Ormdl3* does not alter glucose tolerance, insulin sensitivity, insulin secretion, islet morphology, or cellular ceramide levels on standard chow diet. When challenged with a high fat diet, while *Ormdl3*^β-/-^ mice did not exhibit any alteration in metabolic parameters or islet architecture, lipidomics analysis revealed significantly higher levels of very long chain ceramides in their islets. Taken together, our results reveal that loss of *Ormdl3* alone is not sufficient to impinge upon β-cell function or whole-body glucose and insulin homeostasis, but loss of *Ormdl3* does alter specific sphingolipid levels.

## Introduction

In obesity, chronically elevated levels of circulating free fatty acids contribute to the *de novo* production of cellular lipids, including sphingolipids. Left unchecked, elevated sphingolipid production can lead to the accumulation of sphingolipid species such as ceramide within the cell (1, 2). In pancreatic β-cells, obesity-directed sphingolipid accumulation contributes to β-cell dysfunction through induction of apoptotic, inflammatory, and cellular stress (1, 3, 4).

Orosomucoid-like proteins (ORMDLs) are an ER-resident protein family that inhibits serine palmitoyltransferase (SPT), the rate limiting enzyme in *de novo* sphingolipid biosynthesis (5-7). ORMDL3 is one of three isoforms of this protein family, and genome-wide association studies (GWAS) have identified *Ormdl3* as an obesity-related gene whose expression negatively correlates with body mass index (8). While this suggests that ORMDL3 plays a role in obesity, the role of ORMDL3 in β-cell physiology and pathology remains unknown.

In the current study, we designed a β-cell-specific *Ormdl3* knockout mouse and investigated the consequences of its loss of function under non-stressed (chow or low-fat diet) and high fat diet (HFD) challenged conditions. Our results showed that *Ormdl3* is dispensable for β-cell function regardless of diet. Indeed, β-cell loss of *Ormdl3* did not alter fasting blood glucose, body weight, islet morphology, glucose tolerance, insulin sensitivity, or insulin secretion. Lipidomics of isolated islets identified elevated levels of very long chain ceramides in HFD-challenged knockout animals. Taken together, our results suggest that *Ormdl3* is not required for β-cell function and survival under physiological or surplus nutrition conditions.

## Results

### Chow-fed *Ormdl3*^β-/-^ mice do not exhibit altered β-cell function or glucose homeostasis

To examine the function of ORMDL3 in β-cells, we generated β-cell specific *Ormdl3* knockout mice (KO; *Ormdl3*^β-/-^) by mating mice harboring floxed (exons 2-4) *Ormdl3* (*Ormdl*^fl/fl^) with mice expressing Cre recombinase under the control of the *Ins1* promoter (Ins1tm1.1 Cre) (9). To determine the efficiency of deletion, we performed qPCR analysis in pancreatic islets isolated from 7-week-old *Ormdl3*^β-/-^ mice (Fig. 1A) and found approximately 80% reduction in *Ormdl3* mRNA levels. We also investigated the expression of *Ormdl1* and *Ormdl2* in these islets and found no significant differences in gene expression level relative to wild type (Wt) mice, suggesting that the deletion of *Ormdl3* in β-cells did not trigger compensation by either *Ormd1* or *Ormdl2* at the mRNA level (Fig. 1A).

**Figure 1:**
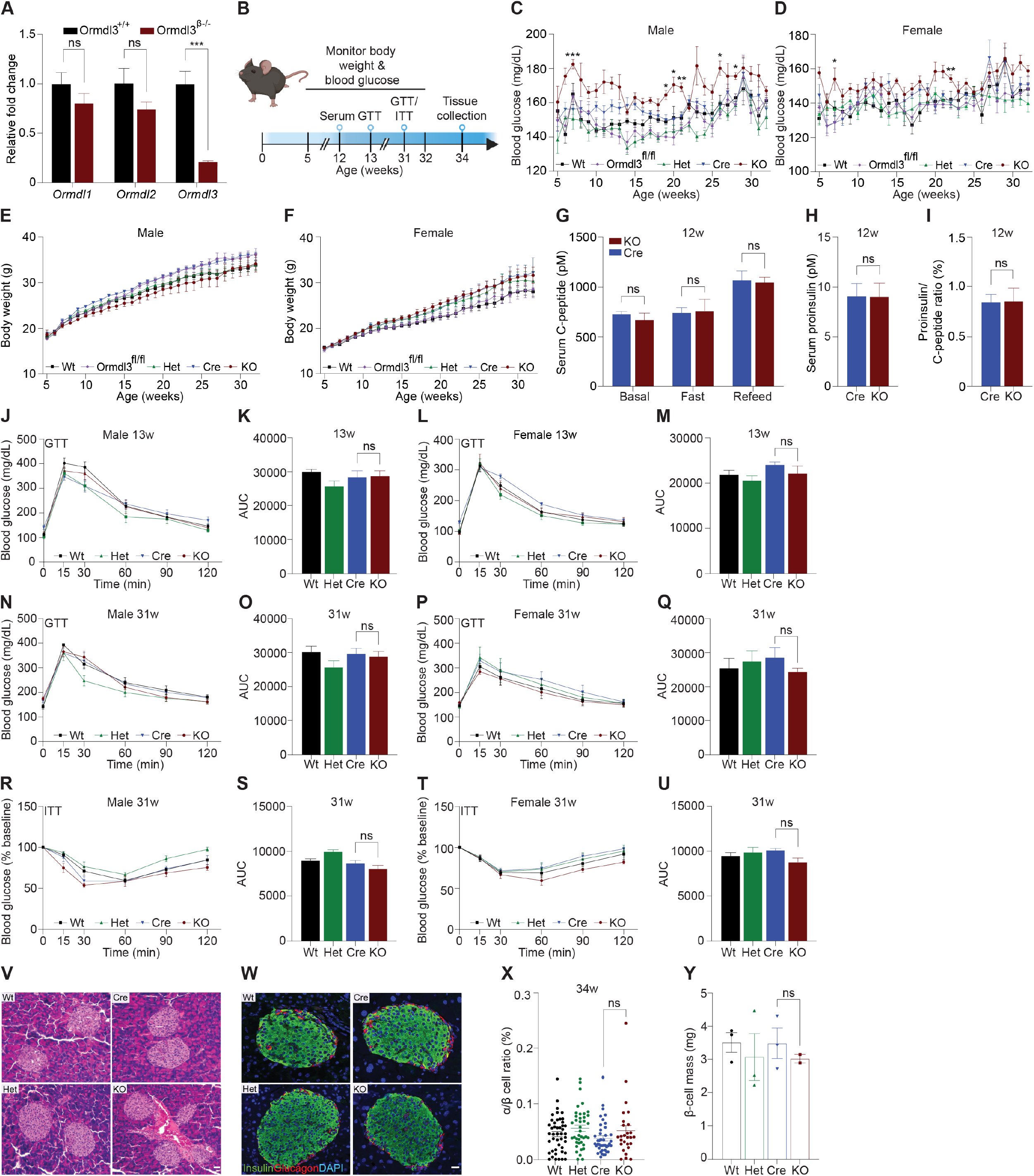
Chow-fed *Ormdl3*^β-/-^ mice do not exhibit altered β-cell function or glucose homeostasis. **A**. Schematic representation of mouse monitoring. **B**. Fold change gene expression of *Ormdl* isoforms in isolated islets of 7-week-old wild-type (Wt) and knockout (KO; *Ormdl3*^β-/-^) mice (n=3/group; Student’s t-test). **C**. Weekly blood glucose measurements in Wt, *Ormdl3*^fl/fl^, Cre, Heterozygous (Het; *Ormdl*^β+/-^), and KO male (n=5-8/group) and **D**. female (n=7-8/group) mice (Two-way ANOVA). **E**. Weekly body weight measurements in Wt, *Ormdl3*^fl/fl^, Cre, Het, and KO male (n=5-8/group) and **F**. female (n=7-8/group) mice (Two-way ANOVA). **G-I**. ELISAs on serum samples from 12-week-old Cre and KO mice (Student’s t-test). **G**. ELISA for C-peptide before/after a 6 hour fast and after 1 hour refeeding (n=5-6/group). **H**. ELISA for proinsulin after 1 hour refeeding (n=5-6/group). **I**. Ratio of proinsulin/C-peptide after 1 hour refeeding (n=5-6/group). **J-Q**. Glucose tolerance and area under the curve (AUC) calculations for male and female Wt, Het, CO, and KO mice. **J**. Glucose tolerance and **K**. total area under the curve (AUC) of 13-week-old male mice (n=5-6/group; Two- and one-way ANOVA, respectively). **L**. Glucose tolerance and **M**. total AUC of 13-week-old female mice (n=5-6/group; Two- and one-way ANOVA, respectively). **N**. Glucose tolerance and **O**. total AUC of 31-week-old male mice (n=5-6/group; Two- and one-way ANOVA, respectively). **P**. Glucose tolerance and **Q**. total AUC of 31-week-old female mice (n=4/group; Two- and one-way ANOVA, respectively). **R-U**. Insulin tolerance and AUC calculations for 31-week-old male and female Wt, Het, Cre, and KO mice **R**. Insulin tolerance and **S**. total AUC of male mice (n=5-6/group; Two- and one-way ANOVA, respectively). **T**. Insulin tolerance and **U**. total AUC of female (n=4/group; Two- and one-way ANOVA, respectively). **V**. Representative images of hematoxylin and eosin staining of 34-week- old male Wt, Cre, Het, and KO mice. **W**. Co-staining for insulin and glucagon in pancreatic sections from 34-week-old male Wt, Cre, Het, and KO mice. **X**. Ratio of α:β cells in co-stained insulin and glucagon sections (n=3/group; One-way ANOVA). **Y**. β-cell mass in insulin-stained sections (n=2-3/group; One-way ANOVA). Data are presented as means ± SEM. One- and two-way ANOVA with Tukey’s post-hoc test (*p<0.05, **p<0.01, ***p<0.001, ****p<0.0001) and Student’s t-test (***p<.001) are applied where indicated. DAPI, 4′,6-diamidino-2-phenylindole; w, weeks; ns, not significant. Scale bars are 20 μm.

We monitored chow-fed male and female Wt, *Ormdl3*^fl/fl^, and Cre mice (as control conditions) as well as <u>*Ormdl3*</u>^β+/-^ (β-cell *Ormdl3* heterozygous knockout; Het), and KO mice starting from 5 weeks of age for 27 weeks (Fig. 1B). During this 27-week period, we measured fasting blood glucose and body weights weekly. We observed a trend towards elevation of blood glucose levels only in male KO mice (Figs. 1C and D) while body weight remained comparable between knockout and control mice (Figs. 1E and F).

To determine if *Ormdl3* ablation would result in impairment of β-cell insulin secretion, we performed fast-refeed experiments in male mice at 12 weeks of age and analyzed serum C-peptide and proinsulin levels (Fig. 1G). Knockout mice did not exhibit altered levels of C-peptide in the *ad libitum* basal condition, after six hours of fasting, or during the refeed condition. Proinsulin levels were also unchanged during the refeed condition (Fig. 1H). The ratio of proinsulin to C-peptide during the refeed condition (Fig. 1I) was unaffected. Taken together, these data suggest that loss of *Ormdl3* in β-cells does not significantly alter insulin secretion.

Next, we examined if the loss-of-function of β-cell *Ormdl3* would result in aberrant glucose homeostasis. We performed glucose and insulin tolerance tests in both male and female mice. Our results showed that there were no significant alterations in glucose tolerance in male and female mice at 13 (Figs. 1J-M) or 31 weeks of age (Figs. 1N-Q). We also measured insulin tolerance in 31-week-old male (Figs. 1R and S) and female mice (Figs. 1T and U) and found no change in insulin sensitivity between KO mice and controls (Wt, Het, and Cre). These results suggest that *Ormdl3* deletion in β-cells does not impair whole-body insulin sensitivity or glucose metabolism.

To determine morphological changes in islets, we performed hematoxylin and eosin (H&E) staining on pancreatic sections obtained from male 34-week-old KO, het, and control (Wt and Cre) mice (Fig. 1V). To investigate if loss of *Ormdl3* alters islet composition and β-cell function, we co-stained sections for insulin and glucagon and analyzed α:β cell ratio (Figs. 1W and X). Our results showed no significant changes in islet morphology (Figs. 1V and W) or β-cell mass (Fig. 1Y). Collectively, our data suggest that under chow-fed conditions, *Ormdl3* deletion is dispensable for β-cell function or glucose homeostasis.

### High fat diet feeding in *Ormdl3*^β-/-^ mice does not impact β-cell function or glucose homeostasis

Our initial results revealed that deletion of *Ormdl3* in the β-cells of mice fed chow diet did not contribute to β-cell dysfunction or impair glucose homeostasis. The ORMDL proteins are known to inhibit serine palmitoyltransferase, the enzyme responsible for the condensation of palmitoyl-CoA and serine into 3-ketosphinganine (Fig. 3A) (10). Since this condensation reaction requires palmitic acid-derived palmitoyl-CoA, we reasoned that low levels of palmitic acid present in chow diet may not lead to sufficient accumulation of sphingolipids underlying β-cell dysfunction. Therefore, we decided to nutritionally challenge these mice with HFD, a model of diet-induced obesity. To this end, we placed male mice on HFD or control low fat diet (LFD) starting at 6 weeks of age for 24 weeks (Fig. 2A) and measured fasting blood glucose levels and body weights weekly. We did not observe any significant changes in fasting blood glucose levels in the LFD-fed Wt, Het, Cre, or KO animals (Fig. 2B). When challenged with a HFD, there were also no changes observed in fasting blood glucose levels (Fig. 2C). Additionally, besides the expected weight gain resulting from high fat diet, there was no significant difference in body weights (Fig. 2D) or food consumption (Figs. 2E and F) between Wt, Het, Cre, or KO mice on either diet.

**Figure 2:**
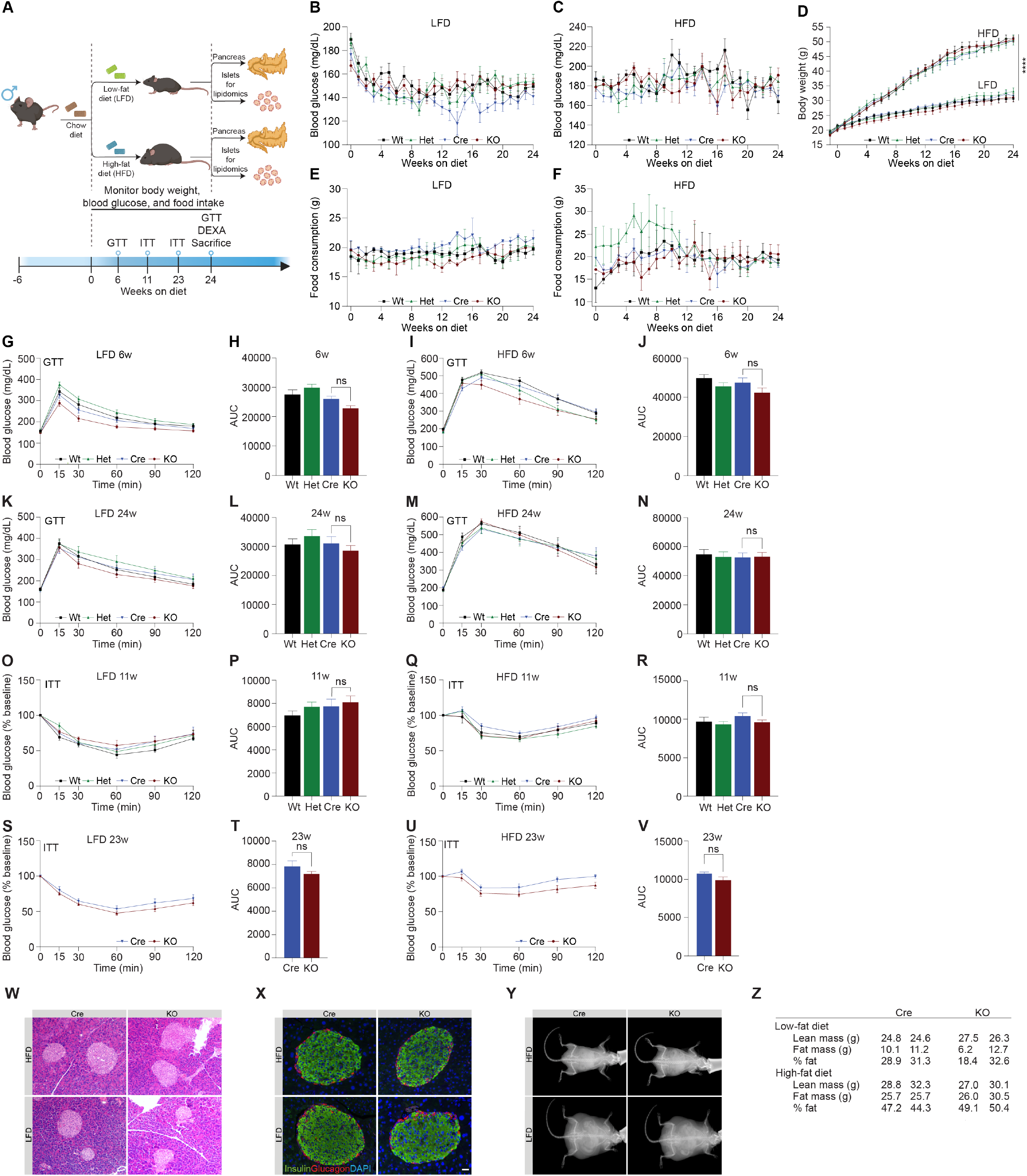
High fat diet feeding in *Ormdl3*^β-/-^ mice does not impact β-cell function or glucose homeostasis. **A**. Schematic representation of mouse monitoring and study design. **B**. Weekly fasting blood glucose assessment of Wt, Het, Cre, and KO mice under LFD and **C**. HFD conditions (n=9/group; two-way ANOVA). **D**. Weekly body weight measurements in LFD and HFD-fed Wt, Het, Cre, and KO mice (n=9/group; Two-way ANOVA). **E**. Weekly food consumption in Wt, Het, Cre, and KO LFD and **F**. HFD-fed mice (grams of food eaten/mouse/week; Two-way ANVOA). **G-N**. Glucose tolerance and area under the curve (AUC) calculations for LFD and HFD-fed male Wt, Het, Cre, and KO mice. **G**. Glucose tolerance and **H**. total AUC of 6-weeks-on-diet LFD-fed mice (n=9/group; Two- and one-way ANOVA, respectively). **I**. Glucose tolerance and **J**. total AUC of 6-weeks-on-diet HFD-fed mice (n=9/group; Two- and one-way ANOVA, respectively). **K**. Glucose tolerance and **L**. total AUC of 24-weeks-on-diet LFD-fed mice (n=9/group; Two- and one-way ANOVA, respectively). **M**. Glucose tolerance and **N**. total AUC of 24-weeks-on-diet HFD-fed mice (n=9/group; Two- and one-way ANOVA, respectively). **O-V**. Insulin tolerance and AUC calculations for LFD and HFD-fed male Wt, Het, Cre, and KO mice. **O**. Insulin tolerance and **P**. total AUC of 11-weeks-on-diet LFD-fed mice (n=9/group; Two- and one-way ANOVA, respectively). **Q**. Insulin tolerance and **R**. total AUC of 11-weeks-on-diet HFD-fed mice (n=9/group; Two- and one-way ANOVA, respectively). **S**. Insulin tolerance and **T**. total AUC of 23-weeks-on-diet LFD-fed mice (n=9/group; Two- and one-way ANOVA, respectively). **U**. Insulin tolerance and **V**. total AUC of 23-weeks-on-diet HFD-fed mice (n=9/group; Two- and one-way ANOVA, respectively). **W**. Representative images of H&E staining of 24-weeks-on-diet male LFD and HFD-fed Cre and KO mice. **X**. Representative images from co-staining for insulin and glucagon in pancreatic sections from 24-weeks-on-diet LFD and HFD-fed male Cre and KO mice. **Y**. Representative images of DEXA assessment in 24-weeks-on-diet LFD and HFD-fed male Cre and KO mice. **Z**. Table of DEXA assessment of lean mass and fat mass (n=2/group).

**Figure 3:**
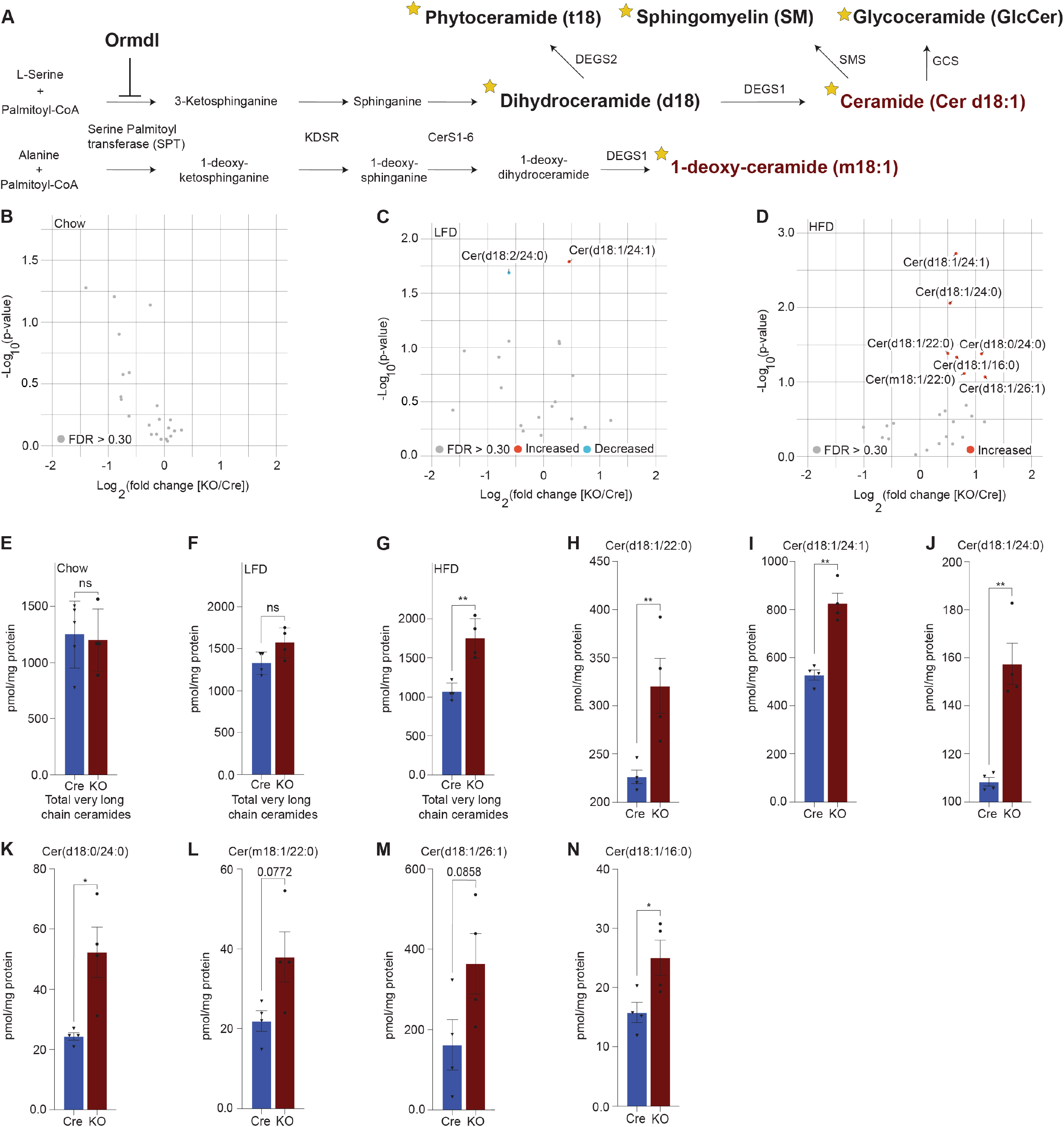
Lipidomics analysis reveals significant upregulation of very long chain ceramide species in islets of high fat diet-fed *Ormdl3*^β-/-^ mice. **A**. Schematic illustration of *de novo* sphingolipid biosynthetic pathway (stars denote lipid classes measured). **B**. Volcano plot of fold change in lipid species between 10-week-old chow-fed Cre and KO mice. **C**. Volcano plot of fold change in lipid species between LFD-fed Cre and KO mice after 24-weeks-on-diet (30 weeks of age). **D**. Volcano plot of fold change in lipid species between HFD-fed Cre and KO mice after 24-weeks-on-diet (30 weeks of age). **E-G**. pmol total very long chain ceramide species (C22-26) normalized to total protein content (mg) in (Welch’s t-test) **E**. 10-week-old chow-fed Cre (n=5) and KO (n=4) mice. **F**. LFD-fed Cre (n=4) and KO (n=4) after 24-weeks-on-diet (30 weeks of age). **G**. HFD-fed Cre (n=4) and KO (n=4) mice after 24-weeks-on-diet (30 weeks of age). **H-N**. pmol ceramide species for HFD-fed mice normalized to total protein content (mg) for (Welch’s t-test) **H**. Cer(d18:1/22:0), **I**. Cer(d18:1/24:1), **J**. Cer(d18:1/24:0), **K**. Cer(d18:0/24:0), **L**. Cer(m18:1/22:0), **M**. Cer(d18:1/26:1), and **N**. Cer(d18:1/16:0). Data are presented as means ± SEM. Unpaired t-test with Welch correction (*p<0.05, **p<0.01) is applied where indicated. mg, milligram; ns, not significant; FDR, false discovery rate; KDSR, 3-keto-dihydrosphingosine reductase; CerS1-6, Ceramide synthase 1-6; DEGS1, Δ4-dihydroceramide desaturase 1; DEGS2, Δ4-dihydroceramide desaturase 2; GCS, glycoceramide synthase; SMS, sphingomyelin synthase. FDR q > .3, grey dots; FDR q < .3, red dots represent upregulated species and blue dots represent downregulated species.

We next examined glucose and insulin tolerance throughout the course of the study. Our results indicated that KO mice did not display any significant difference in glucose tolerance compared to Cre mice in either the LFD or HFD condition at 6 (Figs. 2G-J) or 24 weeks of age (Figs. 2K-N). In addition, we also observed no significant effects of *Ormdl3* deletion on insulin tolerance in LFD or HFD-fed mice at 11 (Figs. 2O-R) and 23 weeks of age (Figs. 2S-V).

To investigate islet morphology, we performed H&E staining of pancreatic sections on LFD and HFD-fed control (Cre) and knockout mice that had been on diet for 24 weeks (Fig. 2W). Further, insulin and glucagon immunofluorescence co-staining showed no significant alterations in islet morphology or composition between HFD and LFD-fed Cre and KO mice (Fig. 2X).

Next, we asked whether loss of *Ormdl3* would result in changes in lean and fat mass (Figs. 2Y and Z). Dual-energy X-ray absorptiometry (DEXA) assessment in 24-week-old LFD and HFD-fed Cre and KO mice showed that there were no significant changes in lean or fat mass in mice on either diet. Taken together, our results suggest that *Ormdl3* deletion does not impinge upon β-cell function and is dispensable for glucose and insulin homeostasis even under obesogenic conditions.

### Lipidomics analysis reveals significant upregulation of very long chain ceramide species in islets of high fat diet-fed *Ormdl3*^β-/-^ mice

Given the inhibitory role of ORMDL proteins in SPT-mediated sphingolipid catabolism (Fig. 3A), we hypothesized that the levels of sphingolipid species downstream of SPT could be altered. To test this, we performed targeted lipidomic analysis (Fig 3A; stars indicate species measured) in isolated islets of chow-fed, LFD-fed, and HFD-fed Cre and *Ormdl3*^β-/*-*^ mice. We first examined fold change between 10-week-old chow-fed *Ormdl3*^β-/-^ and Cre mice, but we did not observe any significantly upregulated or downregulated ceramides species between groups (Fig. 3B). We next analyzed the levels of ceramides from LFD-fed or HFD-challenged *Ormdl3*^β-/*-*^ mice after 24 weeks of feeding (Fig 2A). Initial fold change comparison between LFD-fed *Ormdl3*^β-/-^ and Cre mice revealed upregulation and downregulation of d18:1/24:1 and d18:2/24:0 ceramides, respectively (Fig. 3C). However, fold change comparison between HFD counterparts revealed striking upregulation of islet long chain ceramide species d18:1/16:0 as well as very long chain ceramide species d18:1/24:1, d18:1/24:0, d18:1/22:0, d18:0/24:0, m18:1/22:0, and d18:1/26:1 as compared to Cre control mice (Fig. 3D). There were no significant differences between total very long chain ceramide species (C22-26) in chow-fed (Fig. 3E) and LFD-fed (Fig. 3F) KO mice as compared to control. Interestingly, levels of total very long chain ceramides species were increased in HFD-fed KO mice compared to Cre mice (Fig. 3G). Furthermore, analysis of individual sphingolipid species confirms these changes are indeed observed between HFD-fed *Ormdl3*^β-/-^ mice as compared to Cre control mice (Figs. 3H-N). Taken together, our results suggest that while loss of β-cell *Ormdl3* alone does not impinge upon systemic metabolism or β-cell function in nutritionally unstressed conditions, deletion of *Ormdl3* results in a significant increase in very long chain ceramide species in obesity.

## Discussion

ORMDL3 has been implicated in a variety of disorders including asthma, inflammatory bowel disease, and obesity (8, 11-13). Recently, it was reported that mice with a whole-body knockout of *Ormdl3* when challenged with cold exposure or HFD exhibit impaired regulation of brown adipose tissue thermogenesis, white adipose tissue browning, and insulin resistance (14). Yet the contribution of β-cells to the systemic glucose homeostasis remained unknown. We previously showed that islets from overweight/obese human donors displayed significant downregulation of *ORMDL3* expression compared with islets from lean donors (15). In contrast, *Ormdl3* was substantially upregulated in the islets of leptin-deficient obese (*ob/ob*) mice compared with lean mice (15). *Ormdl3* knockdown in a murine β-cell line induced expression of pro-apoptotic markers suggesting a role for *Ormdl3* in β-cell apoptosis (15). In this study, we found that genetic ablation of *Ormdl3* did not affect glucose metabolism, insulin sensitivity, insulin secretion, or islet architecture. However, despite seemingly unaltered β-cell health and function, targeted lipidomic determination of ceramide levels revealed increases in very long chain ceramides (C22-C26) and long chain C16 ceramide when *Ormdl3*^β-/-^ mice were fed HFD. While our results suggest that β-cell expression of *Ormdl3* is dispensable for normal islet function and glycemic control, they also suggest that ORMDL3 regulates sphingolipid levels in the β-cell.

*Ormdl3* ablation has generated conflicting reports on SPT regulation and sphingolipid metabolism (16, 17). For instance, sphingolipid levels were found to be unchanged in transgenic *Ormdl3* overexpression and whole-body *Ormdl3* KO mice, and this observation was recapitulated in HEK cells (17). However, many studies report some effect of *Ormdl3* overexpression or deletion on systemic sphingolipid levels with minor changes to rodent health measures (14, 18, 19). For example, while absence of *Ormdl3* has been associated with increased levels of sphingolipids within the brain, *Ormdl3* knockout mice appeared metabolically healthy and similar to control mice (19). Our results are in line with these reports demonstrating increased ceramide levels following *Ormdl3* ablation.

Our data indicate that β-cell loss of *Ormdl3* during obesity leads to increased generation of islet very long chain ceramide species, hinting at a potential regulatory axis in which ORMDL3 contributes to control of this sphingolipid class. In mammals, CERS2 catalyzes the N-acylation of the sphingoid base with very long chain fatty acyl-CoAs during *de novo* sphingolipid production to produce very long chain ceramides species (C22-26) (20, 21). Additionally, reports suggest that the ratio of very long chain:long chain ceramides is important for proper cellular function (20, 22). For instance, in BALB/c primary mouse hepatocytes overexpressing *Cers2*, the ratio of very long chain:long chain ceramides was increased, but despite the increase in overall ceramide abundance, insulin signal transduction was improved while markers of ER stress and gluconeogenesis were reduced (22). In addition, a recent report proposes an obesity-independent CERS2-dependent lipid signature of imbalanced very long chain:long chain ceramides as contributing to β-cell failure through impaired proinsulin processing (23). Our findings suggest that very long chain ceramide species may be neither beneficial nor deleterious within the β-cell, but future work including feeding with a ketogenic diet containing a higher fat content could be performed to determine the optimal abundance and ratio of very long acyl chain-containing lipids for normal β-cell health.

Taken together, our findings suggest that while loss-of-function of β-cell ORMDL3 does not impair metabolic health, ORMDL3 contributes to the regulation of very long chain sphingolipid generation in β-cells under obesogenic conditions.

## Materials and Methods

### Animals

The animal care and experimental procedures were carried out in accordance with the recommendations of the National Institutes of Health Guide for the Care and Use of Laboratory Animals. The protocol (#M005064-R01-A02 by F.E. for mice) was approved by the University of Wisconsin-Madison Institutional Animal Care and Use Committee. Mice were bred and maintained under specific pathogen-free conditions at University of Wisconsin-Madison under approved protocols and were housed at 20–24°C on a 12 h light/12 h dark cycle. Animals were observed daily for health status, any mice that met IACUC criteria for euthanasia were immediately euthanized.

### Generation of β-cell specific Ormdl3 knockout mice

β-cell specific *Ormdl3* knockout mice (*Ormdl3*^β-/-^) were generated by Cyagen Biosciences on C57BL/6N background. Briefly, a targeting vector containing mouse *Ormdl3* was generated, containing LoxP sites surrounding exons 2-4 of the *Ormdl3* gene and a Neo selection cassette flanked by Frt motifs. This construct was then electroporated into embryonic stem cells (ESC) of C57BL/6N background, and the resulting cells were then screened for homologous recombination. The ESCs were then validated, and neo-excision was achieved *in vitro* by electroporation with an Flp-O expression plasmid. The resulting neo-excised ESC clones were then screened for successful deletion by PCR and injected into blastocysts isolated from pregnant B6 albino B6(Cg)-Tyr^c-2J^/J females. This resulted in generation of *Ormdl*^fl/fl^ mice on C57BL/6N background. We then bred these mice with commercially available mice expressing a Cre construct expressed under the control of the insulin promotor (B6(Cg)-Ins1^tm1.1(cre)Thor/J^) to delete *Ormdl3* specifically from β-cells.

### Diets, feeding regimen, and weekly measurements

Chow, low fat diet, and high fat diet (Envigo 2919, Research Diets D12450J, and Research diets D12492, respectively) fed animals had *ad libitum* access to food and water unless otherwise specified. Mice that were fed a 60% high fat diet (Research Diets D12492) and 10% low fat diet (Research Diets D12492) began feeding on diet at 6 weeks of age and extending for a period of 24 weeks. Experiments on chow fed mice were performed on male and female mice between 5 and 34 weeks of age. Experiments on low-fat diet and high-fat diet fed mice were performed on male mice between 5 and 30 weeks of age. Weekly assessment of blood glucose and body weights was done after 6 hours of fasting. Blood glucose was analyzed by glucometer (Contour Next EZ 9628) after tail snip.

### Histology

Pancreata from mice were fixed with 10% zinc formalin overnight and paraffin embedded. 5-μm sections of the pancreata were generated, and staining was performed after blocking with 5% normal goat serum with anti-Insulin (Linco) and anti-Glucagon (Cell Signaling) antibodies using established protocols. Antigen retrieval was prepared by using citrate buffer pH 6.0. After staining, slides were mounted with antifade mounting medium containing 4,6-diamidino-2-phenylindole (DAPI) (Vector Laboratories). Immunofluorescent images of pancreatic sections were obtained using a Nikon Storm/Tirf/Epifluorescence. Images used for β-cell mass calculations were obtained with an EVOS FL Auto imaging system. The images of hematoxylin and eosin (H&E)-stained pancreatic sections were obtained using an AmScope light microscope. For analysis of β-cell mass and α:β cell ratio, the images were analyzed by using the Nikon Elements Advanced Research software program.

### Islet isolation

Islets were isolated using the standard collagenase/protease digestion method as previously described (15, 24). Briefly, the pancreatic duct was cannulated and distended with 4°C collagenase/protease solution using Collagenase P (Sigma-Aldrich, USA) in 1x Hank’s balanced salt solution and 0.02% bovine serum albumin. The protease reaction was stopped using RPMI 1640 with 10% fetal bovine serum. Islets were separated from the exocrine tissue using Histopaque-1077 (Sigma-Aldrich, USA). Hand-picked islets were spun briefly at 1000 rpm for 1 minute before snap freezing in liquid nitrogen and storage at -80°C.

### C-peptide and proinsulin ELISA

For measurement of serum C-peptide and proinsulin, blood was collected from mice via tail snip. Whole blood was then spun at 9000 rpm for 7 minutes and serum was collected and stored at - 80°C. Frozen serum was thawed and measured by ELISA for proinsulin (Mercodia, 10-1232-01) and C-peptide (Alpco 80-CPTMS-E01). Samples were analyzed in duplicate.

### Glucose and insulin tolerance tests

Glucose tolerance tests were performed on wild-type, heterozygous, knockout, and Cre control mice on chow, low fat, and high fat diets after a 6-hour morning fast. Blood glucose levels were measured at 0, 15, 30, 60, 90, and 120 minutes after an intraperitoneal administration of glucose at dose of 2g/kg body weight or insulin at dose of .75 U/kg body weight. Blood glucose measurements were measured using a glucometer (Contour Next EZ 9628). Blood glucose readings above the limit of detection were input as 600 mg/dL.

### RT-qPCR

Total RNA was extracted from *Ormdl3*^β+/+^ and *Ormdl3*^β-/-^ mouse islets using TRIzol reagent (Invitrogen) according to manufacturer’s instructions. cDNAs were synthesized from extracted RNA by using Superscript III First Strand RT-PCR kit (Invitrogen). Real-time quantitative PCR amplifications were performed on CFX96 Touch Real-time PCR detection system (Bio-Rad). β-Actin was used as internal control for the quantity of the cDNAs in real time PCR assays.

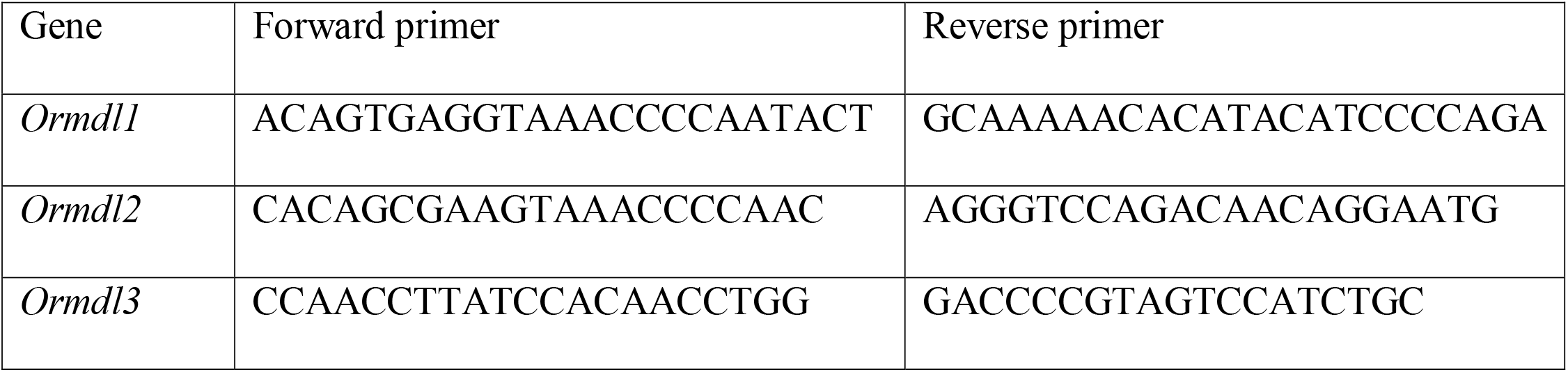
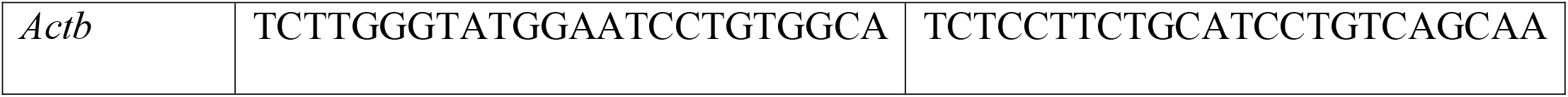

### DEXA measurement

DEXA assessment of lean and fat mass was performed in 24-week-old low fat and high fat diet fed male mice. Mice were anesthetized with isoflurane before determination of body composition using a Faxitron Ultrafoucus in DXA mode. All measurements of body composition were performed in the University of Wisconsin – Madison Small Animal Imaging Facility Core.

### Lipid Extraction

70-90 isolated pancreatic islets were homogenized in 215 μL MeOH with internal standard (final concentrations in supplementary table 1) using a Qiagen TissueLyzer II (9244420) for 4×40s cycles using chilled (4 °C) blocks. 750 μL MTBE was added followed by 250 μL water. Samples were mixed by inversion and phase separation was carried out by centrifugation at 4 °C for 10 min at 16,000 *g*. The top organic phase was transferred to a new 1.5 mL tube and extracts were dried in a SpeedVac. Samples were resuspended in 50:50 ACN/MeOH. A processed blank was prepared in the same way without internal standards added. Samples were stored at -20 °C for 1 week before analysis.

### LC-MS Parameters

LC-MS analysis was performed on an Agilent 1290 Infinity II UHPLC system coupled to an Agilent 6495C triple quadrupole MS. Lipid extracts were separated on an Acquity BEH C18 column (Waters 186009453; 1.7 μm 2.1 × 100 mm) with a VanGuard BEH C18 precolumn (Waters 18003975) maintained at 60°C. Samples were held at 4 °C in a multisampler prior to analysis. Sphingolipids were detected with multiple reaction monitoring (MRM) in positive ion mode. The gas temperature was 210 °C with flow of 11 l/min and the sheath gas temperature was 400 °C with a flow of 12 l/min. The nebulizer pressure was 30 psi, the capillary voltage was set at 4000 V, and the nozzle voltage at 500 V. High pressure RF was 190 and low-pressure RF was 120. Sample injection volume was 10 μL and the injection order was randomized. The chromatography gradient consisted of mobile phase A containing 60:40 ACN/H_2_O in 10 mM ammonium formate and 0.1% formic acid and mobile phase B containing 90:9:1 IPA/ACN/H_2_O in 10 mM ammonium formate and 0.1% formic acid at a flow rate of 0.500 mL/min. The gradient began with 30% B, increasing to 60% over 1.8 min, then increasing to 80% until 7 min, and 99% until 7.14 min held until 10 minutes.

Collision energies, retention times, and scanning windows were optimized based on standards and pooled plasma lipid extracts. Sphingolipid class MRM transitions from are found in supplementary table 1. Retention times for sphingolipids without standards were adjusted using a standard of similar acyl chain length and full scan analysis with a matching chromatography gradient.

### Data Processing

Quantification was performed in the Agilent MassHunter Workstation. Volcano plots and bar graphs were made using the ggpubr package in R version 4.1.2. Data were normalized to protein content as measured by BCA assay (Thermo Scientific 23225) on the pellet in the aqueous phase following lipid extraction. Data is considered “semi-quantitative” because standards were not available for all compounds detected.

## Acknowledgements

We thank Dr. William Holland for providing the protocol and guidance on islet lipidomic studies. H.L. was supported by the NIH National Research Service Award T32 GM007215 and a University of Wisconsin Stem Cell and Regenerative Medicine Center Graduate Fellowship. J.S. is supported by grants from the Juvenile Diabetes Research Foundation (JDRF201309442), the Glenn Foundation for Medical Research (GFMR) and American Federation for Aging Research (AFAR) (22068), the National Institutes of Health (R01DK133479), and the University of Wisconsin-Madison College of Agricultural and Life Science’s Hatch Grant (WIS04000). F.E. is supported by grants from the National Institutes of Health (DK130919 and DK128136), the Juvenile Diabetes Research Foundation (3-SRA-2023-1315-S-B), Greater Milwaukee Foundation, and startup funds from the University of Wisconsin-Madison.

## Author contributions

L.D.H. and H.L. designed and performed the *in vitro and in vivo* experiments, analyzed the data, and prepared the figures. L.D.H. wrote the manuscript. G.W. performed lipidomics analysis and prepared the figures. J.S. interpreted lipidomics data. F.E. conceived, supervised and supported the project, designed experiments, interpreted results and wrote the manuscript. All authors revised the manuscript.

---

**Supplementary Table 1.**

| Cpd Group | Cpd Name | ISTD? | Prec<br>lon | MS1 Res | Prod<br>lon | MS2 Res | Frag<br>(V) | CE<br>(V) | Cell<br>Acc<br>(V) | Ret<br>Time<br>(min) | Ret<br>Window | Polarity |
| --- | --- | --- | --- | --- | --- | --- | --- | --- | --- | --- | --- | --- |
| DHCeramides | Cer(d18:0/02:0) | No | 344.3 | Unit/Enh<br>(6490) | 266.3 | Unit/Enh<br>(6490) | 166 | 30 | 7 | 1.4 | 1 | Positive |
| DHCeramides | Cer(d18:0/02:0) | No | 344.3 | Unit/Enh<br>(6490) | 60.3 | Unit/Enh<br>(6490) | 166 | 28 | 7 | 1.4 | 1 | Positive |
| DHCeramides | Cer(d18:0/06:0) | No | 400.4 | Unit/Enh<br>(6490) | 284.4 | Unit/Enh<br>(6490) | 166 | 30 | 2 | 1.8 | 1 | Positive |
| DHCeramides | Cer(d18:0/06:0) | No | 400.4 | Unit/Enh<br>(6490) | 266.3 | Unit/Enh<br>(6490) | 166 | 30 | 2 | 1.8 | 1 | Positive |
| DHCeramides | Cer(d18:0/08:0) | No | 428.4 | Unit/Enh<br>(6490) | 284.3 | Unit/Enh<br>(6490) | 166 | 24 | 2 | 2 | 1 | Positive |
| DHCeramides | Cer(d18:0/08:0) | No | 428.4 | Unit/Enh<br>(6490) | 266.3 | Unit/Enh<br>(6490) | 166 | 30 | 2 | 2 | 1 | Positive |
| DHCeramides | Cer(d18:0/12:0) | No | 484.5 | Unit/Enh<br>(6490) | 284.3 | Unit/Enh<br>(6490) | 166 | 24 | 7 | 2.4 | 1 | Positive |
| DHCeramides | Cer(d18:0/12:0) | No | 484.5 | Unit/Enh<br>(6490) | 266.3 | Unit/Enh<br>(6490) | 166 | 30 | 7 | 2.4 | 1 | Positive |
| DHCeramides | Cer(d18:0/16:0) | No | 540.5 | Unit/Enh<br>(6490) | 284.3 | Unit/Enh<br>(6490) | 166 | 36 | 5 | 3.3 | 1 | Positive |
| DHCeramides | Cer(d18:0/16:0) | No | 540.5 | Unit/Enh<br>(6490) | 266.3 | Unit/Enh<br>(6490) | 166 | 30 | 4 | 3.3 | 1 | Positive |
| DHCeramides | Cer(d18:0/18:0) | No | 568.6 | Unit/Enh<br>(6490) | 284.3 | Unit/Enh<br>(6490) | 166 | 36 | 6 | 3.8 | 1 | Positive |
| DHCeramides | Cer(d18:0/18:0) | No | 568.6 | Unit/Enh<br>(6490) | 266.3 | Unit/Enh<br>(6490) | 166 | 30 | 6 | 3.8 | 1 | Positive |
| DHCeramides | Cer(d18:0/20:0) | No | 596.6 | Unit/Enh<br>(6490) | 284.4 | Unit/Enh<br>(6490) | 166 | 32 | 3 | 4.2 | 1 | Positive |
| DHCeramides | Cer(d18:0/20:0) | No | 596.6 | Unit/Enh<br>(6490) | 266.3 | Unit/Enh<br>(6490) | 166 | 30 | 3 | 4.2 | 1 | Positive |
| DHCeramides | Cer(d18:0/22:0) | No | 624.6 | Unit/Enh<br>(6490) | 284.4 | Unit/Enh<br>(6490) | 166 | 32 | 3 | 4.6 | 1 | Positive |
| DHCeramides | Cer(d18:0/22:0) | No | 624.6 | Unit/Enh<br>(6490) | 266.3 | Unit/Enh<br>(6490) | 166 | 30 | 3 | 4.6 | 1 | Positive |
| DHCeramides | Cer(d18:0/24:0) | No | 652.7 | Unit/Enh<br>(6490) | 284.4 | Unit/Enh<br>(6490) | 166 | 32 | 3 | 5.5 | 1 | Positive |
| DHCeramides | Cer(d18:0/24:0) | No | 652.7 | Unit/Enh<br>(6490) | 266.3 | Unit/Enh<br>(6490) | 166 | 30 | 3 | 5.5 | 1 | Positive |
| DHCeramides | Cer(d18:0/24:1) | No | 650.7 | Unit/Enh<br>(6490) | 284.3 | Unit/Enh<br>(6490) | 166 | 28 | 3 | 5.2 | 1 | Positive |
| DHCeramides | Cer(d18:0/24:1) | No | 650.7 | Unit/Enh<br>(6490) | 266.3 | Unit/Enh<br>(6490) | 166 | 30 | 3 | 5.2 | 1 | Positive |
| Ceramides | Cer(d18:1/02:0) | No | 342.3 | Unit/Enh<br>(6490) | 264.3 | Unit/Enh<br>(6490) | 166 | 30 | 7 | 1.4 | 1 | Positive |
| Ceramides | Cer(d18:1/02:0) | No | 342.3 | Unit/Enh<br>(6490) | 252.3 | Unit/Enh<br>(6490) | 166 | 30 | 7 | 1.4 | 1 | Positive |
| Ceramides | Cer(d18:1/12:0) | No | 464.5 | Unit/Enh<br>(6490) | 282.3 | Unit/Enh<br>(6490) | 166 | 30 | 4 | 2 | 1 | Positive |
| Ceramides | Cer(d18:1/12:0) | No | 464.5 | Unit/Enh<br>(6490) | 264.2 | Unit/Enh<br>(6490) | 166 | 30 | 4 | 2 | 1 | Positive |
| Ceramides | Cer(d18:1/14:0) | No | 492.5 | Unit/Enh<br>(6490) | 282.3 | Unit/Enh<br>(6490) | 166 | 30 | 4 | 2.7 | 1 | Positive |
| Ceramides | Cer(d18:1/14:0) | No | 492.5 | Unit/Enh<br>(6490) | 264.2 | Unit/Enh<br>(6490) | 166 | 30 | 4 | 2.7 | 1 | Positive |
| Ceramides | Cer(d18:1/16:0) | No | 520.5 | Unit/Enh<br>(6490) | 282.3 | Unit/Enh<br>(6490) | 166 | 30 | 4 | 3.4 | 1 | Positive |
| Ceramides | Cer(d18:1/16:0) | No | 520.5 | Unit/Enh<br>(6490) | 264.2 | Unit/Enh<br>(6490) | 166 | 30 | 4 | 3.4 | 1 | Positive |
| Ceramides | Cer(d18:1/16:1) | No | 518.5 | Unit/Enh<br>(6490) | 282.3 | Unit/Enh<br>(6490) | 166 | 30 | 4 | 2.7 | 1 | Positive |
| Ceramides | Cer(d18:1/16:1) | No | 518.5 | Unit/Enh<br>(6490) | 264.2 | Unit/Enh<br>(6490) | 166 | 30 | 4 | 2.7 | 1 | Positive |
| Ceramides | Cer(d18:1/17:0) | No | 534.5 | Unit/Enh<br>(6490) | 282.3 | Unit/Enh<br>(6490) | 166 | 30 | 4 | 3.5 | 1 | Positive |
| Ceramides | Cer(d18:1/17:0) | No | 534.5 | Unit/Enh<br>(6490) | 264.2 | Unit/Enh<br>(6490) | 166 | 30 | 4 | 3.5 | 1 | Positive |
| Ceramides | Cer(d18:1/18:0) | No | 548.5 | Unit/Enh<br>(6490) | 282.3 | Unit/Enh<br>(6490) | 166 | 30 | 4 | 3.8 | 1 | Positive |
| Ceramides | Cer(d18:1/18:0) | No | 548.5 | Unit/Enh<br>(6490) | 264.2 | Unit/Enh<br>(6490) | 166 | 30 | 4 | 3.8 | 1 | Positive |
| Ceramides | Cer(d18:1/18:1) | No | 546.5 | Unit/Enh<br>(6490) | 282.3 | Unit/Enh<br>(6490) | 166 | 30 | 4 | 3.4 | 1 | Positive |
| Ceramides | Cer(d18:1/18:1) | No | 546.5 | Unit/Enh<br>(6490) | 264.2 | Unit/Enh<br>(6490) | 166 | 30 | 4 | 3.4 | 1 | Positive |
| Ceramides | Cer(d18:1/19:0) | No | 562.6 | Unit/Enh<br>(6490) | 282.3 | Unit/Enh<br>(6490) | 166 | 30 | 4 | 3.4 | 1 | Positive |
| Ceramides | Cer(d18:1/19:0) | No | 562.6 | Unit/Enh<br>(6490) | 264.2 | Unit/Enh<br>(6490) | 166 | 30 | 4 | 3.4 | 1 | Positive |
| Ceramides | Cer(d18:1/20:0) | No | 576.6 | Unit/Enh<br>(6490) | 282.3 | Unit/Enh<br>(6490) | 166 | 30 | 4 | 4.3 | 1 | Positive |
| Ceramides | Cer(d18:1/20:0) | No | 576.6 | Unit/Enh<br>(6490) | 264.2 | Unit/Enh<br>(6490) | 166 | 30 | 4 | 4.3 | 1 | Positive |
| Ceramides | Cer(d18:1/22:0) | No | 604.6 | Unit/Enh<br>(6490) | 282.3 | Unit/Enh<br>(6490) | 166 | 30 | 4 | 4.7 | 1 | Positive |
| Ceramides | Cer(d18:1/22:0) | No | 604.6 | Unit/Enh<br>(6490) | 264.2 | Unit/Enh<br>(6490) | 166 | 30 | 4 | 4.7 | 1 | Positive |
| Ceramides | Cer(d18:1/22:1) | No | 602.6 | Unit/Enh<br>(6490) | 282.3 | Unit/Enh<br>(6490) | 166 | 30 | 4 | 4.2 | 1 | Positive |
| Ceramides | Cer(d18:1/22:1) | No | 602.6 | Unit/Enh<br>(6490) | 264.2 | Unit/Enh<br>(6490) | 166 | 30 | 4 | 4.2 | 1 | Positive |
| Ceramides | Cer(d18:1/23:0) | No | 618.6 | Unit/Enh<br>(6490) | 282.3 | Unit/Enh<br>(6490) | 166 | 30 | 5 | 4.9 | 1 | Positive |
| Ceramides | Cer(d18:1/23:0) | No | 618.6 | Unit/Enh<br>(6490) | 264.2 | Unit/Enh<br>(6490) | 166 | 30 | 4 | 4.9 | 1 | Positive |
| Ceramides | Cer(d18:1/24:0) | No | 632.6 | Unit/Enh<br>(6490) | 282.3 | Unit/Enh<br>(6490) | 166 | 30 | 5 | 5.2 | 1 | Positive |
| Ceramides | Cer(d18:1/24:0) | No | 632.6 | Unit/Enh<br>(6490) | 264.2 | Unit/Enh<br>(6490) | 166 | 30 | 4 | 5.2 | 1 | Positive |
| Ceramides | Cer(d18:1/24:1) | No | 630.6 | Unit/Enh<br>(6490) | 282.3 | Unit/Enh<br>(6490) | 166 | 30 | 4 | 4.7 | 1 | Positive |
| Ceramides | Cer(d18:1/24:1) | No | 630.6 | Unit/Enh<br>(6490) | 264.2 | Unit/Enh<br>(6490) | 166 | 30 | 4 | 4.7 | 1 | Positive |
| Ceramides | Cer(d18:1/26:0) | No | 660.7 | Unit/Enh<br>(6490) | 282.3 | Unit/Enh<br>(6490) | 166 | 30 | 4 | 5.8 | 1 | Positive |
| Ceramides | Cer(d18:1/26:0) | No | 660.7 | Unit/Enh<br>(6490) | 264.2 | Unit/Enh<br>(6490) | 166 | 30 | 4 | 5.8 | 1 | Positive |
| Ceramides | Cer(d18:1/26:1) | No | 658.7 | Unit/Enh<br>(6490) | 282.3 | Unit/Enh<br>(6490) | 166 | 30 | 4 | 5.2 | 1 | Positive |
| Ceramides | Cer(d18:1/26:1) | No | 658.7 | Unit/Enh<br>(6490) | 264.2 | Unit/Enh<br>(6490) | 166 | 30 | 5 | 5.2 | 1 | Positive |
| IS | Cer(d18:1/8:0) | Yes | 426.688 | Unit/Enh<br>(6490) | 282.3 | Unit/Enh<br>(6490) | 166 | 30 | 4 | 2.8 | 1 | Positive |
| IS | Cer(d18:1/8:0) | Yes | 426.688 | Unit/Enh<br>(6490) | 264.2 | Unit/Enh<br>(6490) | 166 | 30 | 4 | 2.8 | 1 | Positive |
| IS | Cer(d18:1d7/15:0) | Yes | 530.917 | Unit/Enh<br>(6490) | 271.3 | Unit/Enh<br>(6490) | 166 | 30 | 4 | 4.1 | 1 | Positive |
| IS | Cer(d18:1d7/16:0) | Yes | 527.5 | Unit/Enh<br>(6490) | 271.3 | Unit/Enh<br>(6490) | 166 | 30 | 4 | 4.3 | 1 | Positive |
| IS | Cer(d18:1d7/18:0) | Yes | 555.5 | Unit/Enh<br>(6490) | 271.3 | Unit/Enh<br>(6490) | 166 | 30 | 4 | 4.8 | 1 | Positive |
| IS | Cer(d18:1d7/24:0) | Yes | 639.6 | Unit/Enh<br>(6490) | 271.3 | Unit/Enh<br>(6490) | 166 | 30 | 4 | 6.4 | 1 | Positive |
| IS | Cer(d18:1d7/24:1) | Yes | 637.6 | Unit/Enh<br>(6490) | 271.3 | Unit/Enh<br>(6490) | 166 | 30 | 4 | 5.8 | 1 | Positive |
| Ceramides | Cer(d18:2/16:0) | No | 536.5 | Unit/Enh<br>(6490) | 280.3 | Unit/Enh<br>(6490) | 166 | 20 | 4 | 3.1 | 1 | Positive |
| Ceramides | Cer(d18:2/16:0) | No | 536.5 | Unit/Enh<br>(6490) | 262.2 | Unit/Enh<br>(6490) | 166 | 20 | 4 | 3.1 | 1 | Positive |
| Ceramides | Cer(d18:2/18:0) | No | 564.5 | Unit/Enh<br>(6490) | 280.3 | Unit/Enh<br>(6490) | 166 | 28 | 4 | 3.6 | 1 | Positive |
| Ceramides | Cer(d18:2/18:0) | No | 564.5 | Unit/Enh<br>(6490) | 262.3 | Unit/Enh<br>(6490) | 166 | 28 | 4 | 3.6 | 1 | Positive |
| Ceramides | Cer(d18:2/20:0) | No | 592.6 | Unit/Enh<br>(6490) | 280.3 | Unit/Enh<br>(6490) | 166 | 28 | 4 | 4.2 | 1 | Positive |
| Ceramides | Cer(d18:2/20:0) | No | 592.6 | Unit/Enh<br>(6490) | 262.3 | Unit/Enh<br>(6490) | 166 | 28 | 4 | 4.2 | 1 | Positive |
| Ceramides | Cer(d18:2/22:0) | No | 620.6 | Unit/Enh<br>(6490) | 280.3 | Unit/Enh<br>(6490) | 166 | 28 | 4 | 5.2 | 1 | Positive |
| Ceramides | Cer(d18:2/22:0) | No | 620.6 | Unit/Enh<br>(6490) | 262.3 | Unit/Enh<br>(6490) | 166 | 28 | 4 | 5.2 | 1 | Positive |
| Ceramides | Cer(d18:2/24:0) | No | 648.6 | Unit/Enh<br>(6490) | 280.3 | Unit/Enh<br>(6490) | 166 | 28 | 4 | 5.7 | 1 | Positive |
| Ceramides | Cer(d18:2/24:0) | No | 648.6 | Unit/Enh<br>(6490) | 262.3 | Unit/Enh<br>(6490) | 166 | 28 | 4 | 5.7 | 1 | Positive |
| Ceramides | Cer(d18:2/24:1) | No | 646.6 | Unit/Enh<br>(6490) | 280.3 | Unit/Enh<br>(6490) | 166 | 28 | 4 | 5 | 1 | Positive |
| Ceramides | Cer(d18:2/24:1) | No | 646.6 | Unit/Enh<br>(6490) | 262.3 | Unit/Enh<br>(6490) | 166 | 28 | 4 | 5 | 1 | Positive |
| deoxyCer | Cer(m18:1/16:0) | No | 522.5 | Unit/Enh<br>(6490) | 266.3 | Unit/Enh<br>(6490) | 166 | 30 | 4 | 3.4 | 1 | Positive |
| deoxyCer | Cer(m18:1/18:0) | No | 550.6 | Unit/Enh<br>(6490) | 266.3 | Unit/Enh<br>(6490) | 166 | 30 | 4 | 3.8 | 1 | Positive |
| deoxyCer | Cer(m18:1/20:0) | No | 578.6 | Unit/Enh<br>(6490) | 266.3 | Unit/Enh<br>(6490) | 166 | 30 | 4 | 4 | 1 | Positive |
| deoxyCer | Cer(m18:1/22:0) | No | 606.6 | Unit/Enh<br>(6490) | 266.3 | Unit/Enh<br>(6490) | 166 | 30 | 4 | 5 | 1 | Positive |
| deoxyCer | Cer(m18:1/24:0) | No | 634.6 | Unit/Enh<br>(6490) | 266.3 | Unit/Enh<br>(6490) | 166 | 30 | 4 | 5.4 | 1 | Positive |
| deoxyCer | Cer(m18:1/24:1) | No | 632.6 | Unit/Enh<br>(6490) | 266.3 | Unit/Enh<br>(6490) | 166 | 30 | 4 | 4.9 | 1 | Positive |
| PhytoCeramides | Cer(t18:0/16:0) | No | 556.5 | Unit/Enh<br>(6490) | 300.3 | Unit/Enh<br>(6490) | 166 | 24 | 4 | 2.6 | 1 | Positive |
| Phytoceramides | Cer(t18:0/16:0) | No | 556.5 | Unit/Enh<br>(6490) | 282.3 | Unit/Enh<br>(6490) | 166 | 24 | 4 | 2.6 | 1 | Positive |
| Phytoceramides | Cer(t18:0/18:0) | No | 584.6 | Unit/Enh<br>(6490) | 300.3 | Unit/Enh<br>(6490) | 166 | 24 | 4 | 3.3 | 1 | Positive |
| Phytoceramides | Cer(t18:0/18:0) | No | 584.6 | Unit/Enh<br>(6490) | 282.4 | Unit/Enh<br>(6490) | 166 | 28 | 4 | 3.3 | 1 | Positive |
| Phytoceramides | Cer(t18:0/20:0) | No | 612.6 | Unit/Enh<br>(6490) | 300.3 | Unit/Enh<br>(6490) | 166 | 24 | 4 | 3.7 | 1 | Positive |
| Phytoceramides | Cer(t18:0/20:0) | No | 612.6 | Unit/Enh<br>(6490) | 282.4 | Unit/Enh<br>(6490) | 166 | 28 | 4 | 3.7 | 1 | Positive |
| Phytoceramides | Cer(t18:0/22:0) | No | 640.6 | Unit/Enh<br>(6490) | 300.3 | Unit/Enh<br>(6490) | 166 | 24 | 4 | 3.7 | 1 | Positive |
| Phytoceramides | Cer(t18:0/22:0) | No | 640.6 | Unit/Enh<br>(6490) | 282.4 | Unit/Enh<br>(6490) | 166 | 28 | 4 | 3.7 | 1 | Positive |
| Phytoceramides | Cer(t18:0/24:0) | No | 668.7 | Unit/Enh<br>(6490) | 300.3 | Unit/Enh<br>(6490) | 166 | 24 | 4 | 4.2 | 1 | Positive |
| Phytoceramides | Cer(t18:0/24:0) | No | 668.7 | Unit/Enh<br>(6490) | 282.4 | Unit/Enh<br>(6490) | 166 | 28 | 4 | 4.2 | 1 | Positive |
| Phytoceramides | Cer(t18:0/24:1) | No | 666.6 | Unit/Enh<br>(6490) | 300.3 | Unit/Enh<br>(6490) | 166 | 24 | 4 | 4 | 1 | Positive |
| Phytoceramides | Cer(t18:0/24:1) | No | 666.6 | Unit/Enh<br>(6490) | 282.4 | Unit/Enh<br>(6490) | 166 | 28 | 4 | 4 | 1 | Positive |
| IS | Cer(d18:0/18:1) | Yes | 566.5 | Unit/Enh<br>(6490) | 284.3 | Unit/Enh<br>(6490) | 166 | 32 | 5 | 4.5 | 1 | Positive |
| IS | Cer(d18:0/18:1) | Yes | 566.5 | Unit/Enh<br>(6490) | 266.3 | Unit/Enh<br>(6490) | 166 | 32 | 5 | 4.5 | 1 | Positive |
| DHGSC | GlcCer<br>(d18:0/16:0) | No | 702.6 | Unit/Enh<br>(6490) | 284.3 | Unit/Enh<br>(6490) | 166 | 29 | 4 | 3.5 | 1 | Positive |
| DHGSC | GlcCer<br>(d18:0/16:0) | No | 702.6 | Unit/Enh<br>(6490) | 266.3 | Unit/Enh<br>(6490) | 166 | 29 | 4 | 3.5 | 1 | Positive |
| DHGSC | GlcCer<br>(d18:0/18:0) | No | 730.6 | Unit/Enh<br>(6490) | 284.3 | Unit/Enh<br>(6490) | 166 | 29 | 4 | 3.3 | 1 | Positive |
| DHGSC | GlcCer<br>(d18:0/18:0) | No | 730.6 | Unit/Enh<br>(6490) | 266.3 | Unit/Enh<br>(6490) | 166 | 29 | 4 | 3.3 | 1 | Positive |
| GSC | GlcCer<br>(d18:0/2:0) | No | 506.4 | Unit/Enh<br>(6490) | 266.3 | Unit/Enh<br>(6490) | 166 | 29 | 4 | 2 | 1 | Positive |
| DHGSC | GlcCer<br>(d18:0/20:0) | No | 758.6 | Unit/Enh<br>(6490) | 284.3 | Unit/Enh<br>(6490) | 166 | 29 | 4 | 4.3 | 1 | Positive |
| DHGSC | GlcCer<br>(d18:0/20:0) | No | 758.6 | Unit/Enh<br>(6490) | 266.3 | Unit/Enh<br>(6490) | 166 | 29 | 4 | 4.3 | 1 | Positive |
| DHGSC | GlcCer<br>(d18:0/22:0) | No | 786.7 | Unit/Enh<br>(6490) | 284.3 | Unit/Enh<br>(6490) | 166 | 29 | 4 | 4.7 | 1 | Positive |
| DHGSC | GlcCer<br>(d18:0/22:0) | No | 786.7 | Unit/Enh<br>(6490) | 266.3 | Unit/Enh<br>(6490) | 166 | 29 | 4 | 4.7 | 1 | Positive |
| DHGSC | GlcCer<br>(d18:0/24:0) | No | 814.7 | Unit/Enh<br>(6490) | 284.3 | Unit/Enh<br>(6490) | 166 | 29 | 4 | 5 | 1 | Positive |
| DHGSC | GlcCer<br>(d18:0/24:0) | No | 814.7 | Unit/Enh<br>(6490) | 266.3 | Unit/Enh<br>(6490) | 166 | 29 | 4 | 5 | 1 | Positive |
| DHGSC | GlcCer<br>(d18:0/24:1) | No | 812.7 | Unit/Enh<br>(6490) | 284.3 | Unit/Enh<br>(6490) | 166 | 29 | 4 | 4.9 | 1 | Positive |
| DHGSC | GlcCer<br>(d18:0/24:1) | No | 812.7 | Unit/Enh<br>(6490) | 266.3 | Unit/Enh<br>(6490) | 166 | 29 | 4 | 4.9 | 1 | Positive |
| GSC | GlcCer<br>(d18:1/16:0) | No | 700.6 | Unit/Enh<br>(6490) | 282.3 | Unit/Enh<br>(6490) | 166 | 29 | 4 | 3.4 | 1 | Positive |
| GSC | GlcCer<br>(d18:1/16:0) | No | 700.6 | Unit/Enh<br>(6490) | 264.3 | Unit/Enh<br>(6490) | 166 | 29 | 4 | 3.4 | 1 | Positive |
| GSC | GlcCer<br>(d18:1/18:0) | No | 728.6 | Unit/Enh<br>(6490) | 282.3 | Unit/Enh<br>(6490) | 166 | 29 | 4 | 3.8 | 1 | Positive |
| GSC | GlcCer<br>(d18:1/18:0) | No | 728.6 | Unit/Enh<br>(6490) | 264.2 | Unit/Enh<br>(6490) | 166 | 29 | 4 | 3.8 | 1 | Positive |
| GSC | GlcCer<br>(d18:1/2:0) | No | 504.4 | Unit/Enh<br>(6490) | 264.3 | Unit/Enh<br>(6490) | 166 | 29 | 4 | 2.1 | 1 | Positive |
| GSC | GlcCer<br>(d18:1/2:0) | No | 504.4 | Unit/Enh<br>(6490) | 174.1 | Unit/Enh<br>(6490) | 166 | 29 | 4 | 2.1 | 1 | Positive |
| GSC | GlcCer<br>(d18:1/20:0) | No | 756.6 | Unit/Enh<br>(6490) | 282.3 | Unit/Enh<br>(6490) | 166 | 29 | 4 | 4.2 | 1 | Positive |
| GSC | GlcCer<br>(d18:1/20:0) | No | 756.6 | Unit/Enh<br>(6490) | 264.2 | Unit/Enh<br>(6490) | 166 | 29 | 4 | 4.2 | 1 | Positive |
| GSC | GlcCer<br>(d18:1/22:0) | No | 784.7 | Unit/Enh<br>(6490) | 282.3 | Unit/Enh<br>(6490) | 166 | 29 | 4 | 4.7 | 1 | Positive |
| GSC | GlcCer<br>(d18:1/22:0) | No | 784.7 | Unit/Enh<br>(6490) | 264.2 | Unit/Enh<br>(6490) | 166 | 29 | 4 | 4.7 | 1 | Positive |
| GSC | GlcCer<br>(d18:1/24:0) | No | 812.7 | Unit/Enh<br>(6490) | 282.3 | Unit/Enh<br>(6490) | 166 | 29 | 4 | 4.6 | 1 | Positive |
| GSC | GlcCer<br>(d18:1/24:0) | No | 812.7 | Unit/Enh<br>(6490) | 264.2 | Unit/Enh<br>(6490) | 166 | 29 | 4 | 4.6 | 1 | Positive |
| GSC | GlcCer<br>(d18:1/24:1) | No | 810.7 | Unit/Enh<br>(6490) | 282.3 | Unit/Enh<br>(6490) | 166 | 29 | 4 | 4.7 | 1 | Positive |
| GSC | GlcCer<br>(d18:1/24:1) | No | 810.7 | Unit/Enh<br>(6490) | 264.2 | Unit/Enh<br>(6490) | 166 | 29 | 4 | 4.7 | 1 | Positive |
| IS | GlcCer<br>(d18:1/8:0) | Yes | 589.44 | Unit/Enh<br>(6490) | 282.3 | Unit/Enh<br>(6490) | 166 | 29 | 4 | 2.6 | 1 | Positive |
| IS | GlcCer<br>(d18:1/8:0) | Yes | 589.44 | Unit/Enh<br>(6490) | 264.3 | Unit/Enh<br>(6490) | 166 | 29 | 4 | 2.6 | 1 | Positive |
| Sphingolipids | S1P | No | 380.3 | Unit/Enh<br>(6490) | 264.2 | Unit/Enh<br>(6490) | 166 | 12 | 4 | 2.6 | 1 | Positive |
| Sphingolipids | S1P | No | 380.3 | Unit/Enh<br>(6490) | 247.2 | Unit/Enh<br>(6490) | 166 | 12 | 4 | 2.6 | 1 | Positive |
| Sphingolipids | S1P | No | 380.3 | Unit/Enh<br>(6490) | 82.1 | Unit/Enh<br>(6490) | 166 | 37 | 4 | 2.6 | 1 | Positive |
| Sphingolipids | Sa1P | No | 382.3 | Unit/Enh<br>(6490) | 284.3 | Unit/Enh<br>(6490) | 166 | 15 | 4 | 3.6 | 1 | Positive |
| Sphingolipids | Sa1P | No | 382.3 | Unit/Enh<br>(6490) | 266.2 | Unit/Enh<br>(6490) | 166 | 20 | 4 | 3.6 | 1 | Positive |
| Sphingolipids | Sa1P | No | 382.3 | Unit/Enh<br>(6490) | 249.2 | Unit/Enh<br>(6490) | 166 | 37 | 4 | 3.6 | 1 | Positive |
| DHSM | SM(d18:0/16:0) | No | 705.6 | Unit/Enh<br>(6490) | 266.3 | Unit/Enh<br>(6490) | 166 | 29 | 4 | 2.6 | 1 | Positive |
| DHSM | SM(d18:0/16:0) | No | 705.6 | Unit/Enh<br>(6490) | 184.4 | Unit/Enh<br>(6490) | 166 | 29 | 4 | 2.6 | 1 | Positive |
| DHSM | SM(d18:0/18:0) | No | 733.6 | Unit/Enh<br>(6490) | 266.3 | Unit/Enh<br>(6490) | 166 | 29 | 4 | 2.6 | 1 | Positive |
| DHSM | SM(d18:0/18:0) | No | 733.6 | Unit/Enh<br>(6490) | 184.4 | Unit/Enh<br>(6490) | 166 | 29 | 4 | 2.6 | 1 | Positive |
| DHSM | SM(d18:0/2:0) | No | 509.4 | Unit/Enh<br>(6490) | 184.1 | Unit/Enh<br>(6490) | 166 | 29 | 4 | 1.7 | 1 | Positive |
| DHSM | SM(d18:0/20:0) | No | 761.6 | Unit/Enh<br>(6490) | 266.3 | Unit/Enh<br>(6490) | 166 | 29 | 4 | 3.9 | 1 | Positive |
| DHSM | SM(d18:0/20:0) | No | 761.6 | Unit/Enh<br>(6490) | 184.4 | Unit/Enh<br>(6490) | 166 | 29 | 4 | 3.9 | 1 | Positive |
| DHSM | SM(d18:0/22:0) | No | 789.7 | Unit/Enh<br>(6490) | 266.3 | Unit/Enh<br>(6490) | 166 | 29 | 4 | 4.4 | 1 | Positive |
| DHSM | SM(d18:0/22:0) | No | 789.7 | Unit/Enh<br>(6490) | 184.4 | Unit/Enh<br>(6490) | 166 | 29 | 4 | 4.4 | 1 | Positive |
| DHSM | SM(d18:0/24:0) | No | 817.7 | Unit/Enh<br>(6490) | 266.3 | Unit/Enh<br>(6490) | 166 | 29 | 4 | 4.8 | 1 | Positive |
| DHSM | SM(d18:0/24:0) | No | 817.7 | Unit/Enh<br>(6490) | 184.4 | Unit/Enh<br>(6490) | 166 | 29 | 4 | 4.8 | 1 | Positive |
| DHSM | SM(d18:0/24:1) | No | 815.7 | Unit/Enh<br>(6490) | 266.3 | Unit/Enh<br>(6490) | 166 | 29 | 4 | 4.4 | 1 | Positive |
| DHSM | SM(d18:0/24:1) | No | 815.7 | Unit/Enh<br>(6490) | 184.4 | Unit/Enh<br>(6490) | 166 | 29 | 4 | 4.4 | 1 | Positive |
| SM | SM(d18:1/14:0) | No | 675.5 | Unit/Enh<br>(6490) | 264.3 | Unit/Enh<br>(6490) | 166 | 29 | 4 | 2.3 | 1 | Positive |
| SM | SM(d18:1/14:0) | No | 675.5 | Unit/Enh<br>(6490) | 184.4 | Unit/Enh<br>(6490) | 166 | 29 | 4 | 2.3 | 1 | Positive |
| SM | SM(d18:1/16:0) | No | 703.6 | Unit/Enh<br>(6490) | 264.3 | Unit/Enh<br>(6490) | 166 | 29 | 4 | 2.5 | 1 | Positive |
| SM | SM(d18:1/16:0) | No | 703.6 | Unit/Enh<br>(6490) | 184.4 | Unit/Enh<br>(6490) | 166 | 29 | 4 | 2.5 | 1 | Positive |
| IS | SM(d18:1/17:0) | Yes | 717.6 | Unit/Enh<br>(6490) | 264.3 | Unit/Enh<br>(6490) | 166 | 29 | 4 | 3 | 1 | Positive |
| IS | SM(d18:1/17:0) | Yes | 717.6 | Unit/Enh<br>(6490) | 184.4 | Unit/Enh<br>(6490) | 166 | 29 | 4 | 3 | 1 | Positive |
| SM | SM(d18:1/18:0) | No | 731.6 | Unit/Enh<br>(6490) | 264.3 | Unit/Enh<br>(6490) | 166 | 29 | 4 | 2.9 | 1 | Positive |
| SM | SM(d18:1/18:0) | No | 731.6 | Unit/Enh<br>(6490) | 184.4 | Unit/Enh<br>(6490) | 166 | 29 | 4 | 2.9 | 1 | Positive |
| IS | SM(d18:1/18:1)d9 | Yes | 738.6 | Unit/Enh<br>(6490) | 264.3 | Unit/Enh<br>(6490) | 166 | 29 | 4 | 3.9 | 1 | Positive |
| IS | SM(d18:1/18:1)d9 | Yes | 738.6 | Unit/Enh<br>(6490) | 184.1 | Unit/Enh<br>(6490) | 166 | 29 | 4 | 3.9 | 1 | Positive |
| SM | SM(d18:1/2:0) | No | 507.4 | Unit/Enh<br>(6490) | 184.1 | Unit/Enh<br>(6490) | 166 | 29 | 4 | 2 | 1 | Positive |
| SM | SM(d18:1/20:0) | No | 759.6 | Unit/Enh<br>(6490) | 264.3 | Unit/Enh<br>(6490) | 166 | 29 | 4 | 3.8 | 1 | Positive |
| SM | SM(d18:1/20:0) | No | 759.6 | Unit/Enh<br>(6490) | 184.4 | Unit/Enh<br>(6490) | 166 | 29 | 4 | 3.8 | 1 | Positive |
| SM | SM(d18:1/22:0) | No | 787.7 | Unit/Enh<br>(6490) | 264.3 | Unit/Enh<br>(6490) | 166 | 29 | 4 | 3.9 | 1 | Positive |
| SM | SM(d18:1/22:0) | No | 787.7 | Unit/Enh<br>(6490) | 184.4 | Unit/Enh<br>(6490) | 166 | 29 | 4 | 3.9 | 1 | Positive |
| SM | SM(d18:1/24:0) | No | 815.7 | Unit/Enh<br>(6490) | 264.3 | Unit/Enh<br>(6490) | 166 | 29 | 4 | 4.5 | 1 | Positive |
| SM | SM(d18:1/24:0) | No | 815.7 | Unit/Enh<br>(6490) | 184.4 | Unit/Enh<br>(6490) | 166 | 29 | 4 | 4.5 | 1 | Positive |
| SM | SM(d18:1/24:1) | No | 813.7 | Unit/Enh<br>(6490) | 264.3 | Unit/Enh<br>(6490) | 166 | 29 | 4 | 4 | 1 | Positive |
| SM | SM(d18:1/24:1) | No | 813.7 | Unit/Enh<br>(6490) | 184.4 | Unit/Enh<br>(6490) | 166 | 29 | 4 | 4 | 1 | Positive |
| Sphinganine | Sphinganine | No | 302.3 | Unit/Enh<br>(6490) | 284.2 | Unit/Enh<br>(6490) | 166 | 8 | 7 | 1.1 | 1 | Positive |
| Sphinganine | Sphinganine | No | 302.3 | Unit/Enh<br>(6490) | 254.3 | Unit/Enh<br>(6490) | 166 | 20 | 7 | 1.1 | 1 | Positive |
| Sphingosine | Sphingosine | No | 300.3 | Unit/Enh<br>(6490) | 282.3 | Unit/Enh<br>(6490) | 166 | 8 | 6 | 1 | 1 | Positive |
| Sphingosine | Sphingosine | No | 300.3 | Unit/Enh<br>(6490) | 252.3 | Unit/Enh<br>(6490) | 166 | 20 | 6 | 1 | 1 | Positive |

